# Anti-inflammatory properties of natural ingredients used in combinations on adjuvant induced arthritis in rats

**DOI:** 10.1101/325365

**Authors:** Amélie St-Pierre, Véronique Desrosiers, France Dufresne, Pierre U. Blier

**Author notes:** Abbreviations: BS, *Boswellia serrata*; SA, *Salix alba*; GS, Glucosamine sulfate; HA, Hyaluronic acid; RA, Rheumatoid arthritis; NSAID, nonsteroidal anti-inflammatory drugs; ROS, Reactive oxygen specie; SCW, Streptococcal cell wall; CIA, Collagen-induced arthritis; CFA, Complete Freund adjuvant.

## Abstract

Background: Rheumatoid arthritis has seen a significant increase in both incidence and prevalence and its treatments show limited efficiency due to their undesirable effects on patient health. Therefore, major interests lie in the development of treatments with drugs derived from plants or other natural sources with little adverse effects as an alternative to current treatments.

Hypothesis/Purpose: The present study evaluates the therapeutic effects of glucosamine against rheumatoid arthritis in combination with hyaluronic acid, resin extract of *Boswellia serrata* or a bark extract of *Salix alba* on an animal model. We suggest that combinations with plants could improve the attenuation of arthritis symptoms and articular inflammation.

Study design: We used Freund’s complete adjuvant on rats as models of rheumatoid arthritis. Individuals were separated into eight experimental groups: a control group without arthritis, one with arthritis and without treatment, and six other groups receiving a daily therapeutic treatment from days 14 to 29.

Methods: Hind-paw thickness and arthritis scores were measured at days 0, 3, 6 and 9 post-induction, and then every day from days 12 to 29 with a digital caliper and a score system respectively. At the end of the treatment, the mRNA content of three pro-inflammatory cytokines from cartilage was measured using real-time PCR. The total antioxidant activity was evaluated with an Antioxidant Assay Kit.

Results: Treatments with *Boswellia serrata* and *Salix alba* (*Glu+Hyal A+Bosw, Glu+Bosw+Sal, Glu+Bosw* and *Glu+Hyal A+Sal*) saw significant reductions in hind-paw thickness and arthritis scores at the end of the experiment when compared to the untreated group. Expression of pro-inflammatory gene *IL 17A* was also reduced, but only the *Glu+Hyal A+Sal* combination significantly decreased the expression of *IL-1β* and *TNF-α*. The total antioxidant activity in blood plasma significantly increased in groups treated with plant extracts.

Conclusion: The addition of *Boswellia serrata* and/or *Salix alba* attenuates clinical signs of rheumatoid arthritis in Freund’s complete adjuvant-induced arthritis in rats likely due to both their anti-inflammatory and antioxidant properties.

## Introduction

Rheumatoid arthritis is an inflammatory autoimmune disorder. Its incidence and prevalence increase considerably all over the world. In North America and North-European countries, its incidence varies between 20 and 50 per 100,000 population (Fazal et al., 2018). It causes inflammation in joints that leads to pain, stiffness, swelling and cartilage damage (Arthritis Society, 2018). Current treatments do not usually regenerate damaged cartilage or slow the degeneration, but relieve symptoms instead (Arthritis Society, 2018). Treatments using steroids, nonsteroidal anti-inflammatory drugs (NSAIDs), topical anti-inflammatories, biological agents (TNF-α and IL-1 antagonists), acetaminophen and injection of corticosteroids and hyaluronic acid are used against joint diseases but show limited efficiency due to their undesirable adverse effects on patient health (Fan et al., 2005; Zheng et al., 2014). These pharmaceutical drugs can provoke gastrointestinal disturbances (ulcers and perforations), cardiovascular complications, reproductive toxicity, loss of bone mass, and topical applications can be of no benefit when the target joints are too deep (Fan et al., 2005; Umar et al., 2014; Zheng et al., 2014). Due to these limitations, there is an important incentive for the development of biomolecules derived from plants or natural sources without adverse effects as an alternative to NSAIDs and other treatments. Among these biomolecules, glucosamine sulfate (GS), hyaluronic acid (HA), resin extracts of indian frankincense (*Boswellia serrata* Roxb. Ex Colebr., BS) and bark extracts of white willow (*Salix alba* L., SA) are four natural ingredients that are individually considered as efficient against arthritis by regulatory agencies (for example see the monograph of Natural and Non-prescription Health Products Directorate (NNHPD) in Canada (Health Canada, 2018)).

GS is an important component of cartilage and is naturally synthesized in the body. This amino monosaccharide stimulates the biosynthesis of glycosaminoglycan chains, giving the cartilage its strength, flexibility and elasticity, all the while possessing anti-inflammatory properties (Singh et al., 2007). HA is a large viscoelastic glycosaminoglycan present in the synovial fluid, and is responsible for its viscoelastic properties (Moreland, 2003). It also confers good protective properties including shock absorption, protective coating of the articular cartilage surface, and lubrication. BS and SA have both anti-inflammatory and analgesic properties due to the presence of boswellic acid and salicin respectively (Kimmatkar et al., 2003; Shara and Stohs, 2015). Boswellic acid reduces pain and swelling, has antioxidant and free radical-scavenging properties, and appears as a potential new treatment of inflammatory disorders like rheumatoid arthritis and osteoarthritis (Umar et al., 2014). It reduces glycosaminoglycan degradation, keeping the cartilage in good condition unlike NSAIDs that can induce the disruption of the glycosaminoglycan synthesis, accelerating articular damage (Kimmatkar et al., 2003). Salicin has anti-inflammatory and anabolic effects, as shown in canine joints (Shara and Stohs, 2015). Benefits of these natural ingredients have so far only been studied separately, and their potential synergistic effects need to be assessed, as a combination of ingredients can improve their therapeutic effects at the low doses recommended by the health regulation agencies.

Thus, the aim of this study is to examine the potency of different combinations of natural ingredients to limit arthritis symptoms and articular inflammation on an animal model of rheumatoid arthritis. In this context, we used rats previously injected with Freund’s complete adjuvant. Therefore, quantifying the modulation of inflammation might represent the extent to which hyaluronic acid, *Boswellia serrata* or *Salix alba* extract combined to glucosamine can improve the therapeutic efficiency of glucosamine alone.

## Materials and methods

### Animals

Adult female Lewis rats (10 weeks old) were obtained from Charles River Laboratories (Montreal, QC, Canada). Animals were kept at Université du Québec à Rimouski (UQAR) in controlled experimental conditions (23 ± 1°C, relative humidity 40–60%, 12h light/dark cycles, water and LabDiet 5002 *ad libitum*). They were acclimated during 1 week before the experiment. Animal manipulation was conducted in accordance with the Institutional Animal Care Committee of Université du Québec à Rimouski (protocol #CPA-66-16-178).

### Adjuvant Induction

Arthritis was induced by subcutaneous injection of 60 μl of Freund’s adjuvant, a solution of *Mycobacterium tuberculosis* inactivated by heat (Chondrex, Inc. Redmond, WA, USA, 10 mg/ml), at the base of the tail. First symptoms of arthritis appeared 12 days after induction.

### Evaluation of Clinical Signs of Arthritis

Arthritis symptoms were examined at days 0, 3, 6 and 9 post-induction, and then every day from days 12 to 29. Hind-paw thickness was measured with a digital caliper. Arthritis scores were determined by a score system: for each of hind paw, a scale of 0–4; 0, no macroscopic sign; 1, irritation (swelling and redness) at one joint; 2, irritation at more than one joint and/or ankles; 3, irritation at many joints and moderate swelling at the ankle; 4, irritation at many joints and severe swelling at the ankle. For each forepaw, a scale of 0–3 was used; 0, no macroscopic sign; 1, irritation at one joint; 2, irritation at many joints and/or wrist; 3, irritation at all joints and moderate to severe swelling at the wrist. The final score was calculated by adding the individual score of each paw for a maximal result of 14 (Aghazadeh-Habashi et al., 2014).

### Therapeutic Ingredient Administration

Rats were separated randomly in eight different groups: a control group without arthritis (control), another one with arthritis and no treatment (CFA, Freund’s complete adjuvant) and six other groups receiving a daily therapeutic treatment from days 14 to 29. The temporal experimentation plan of animal manipulations is presented in Fig 1. Six therapeutic treatments were administered to the rats: GS (*Glu*); GS and HA (*Glu+Hyal A*); GS, HA and BS (*Glu+Hyal A+Bosw*); GS, HA and SA (*Glu+Hyal A+Sal*); GS and BS *(Glu+Bosw*); and GS, BS and SA (*Glu+Bosw+Sal*).

**Fig 1.**
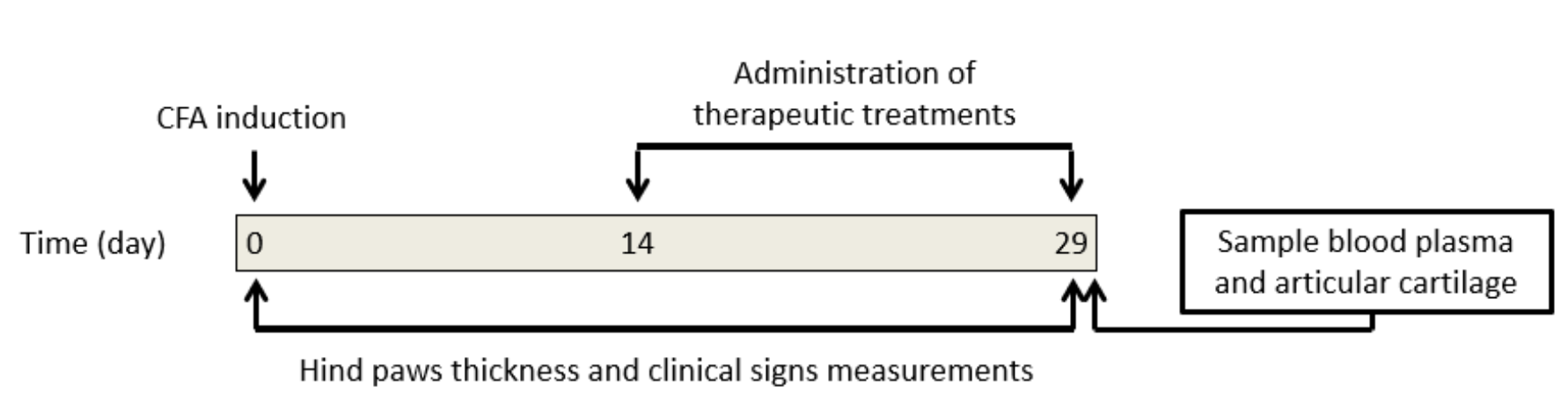
Temporal experimental plan of animal manipulations.

Each ingredient was administered individually, even when the therapeutic treatment had many compounds, and orally (back of the mouth) with a pipette. For each ingredient, the daily dose corresponded to the maximal recommended dosage for humans by Natural and Non-prescription Health Products Directorate (NNHPD) of Health Canada (2014) in its monograph titled «Multiple ingredient joint health products». The dosages for rats were calculated considering an average human weight of 60 kg (weight approved by Food and Drugs Administration (FDA) for safety studies) and a normalization that takes into account the body surface (Reagan-Shaw et al., 2007). The daily dosages are presented in Table 1. This formula represents the allometric conversion:

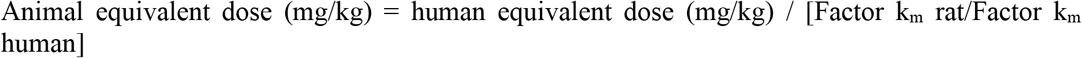

where

Factor k_m_ = body mass (kg) / total body surface (m^2^)

and

k_m_ rat = 6
k_m_ human = 37

**Table 1.**
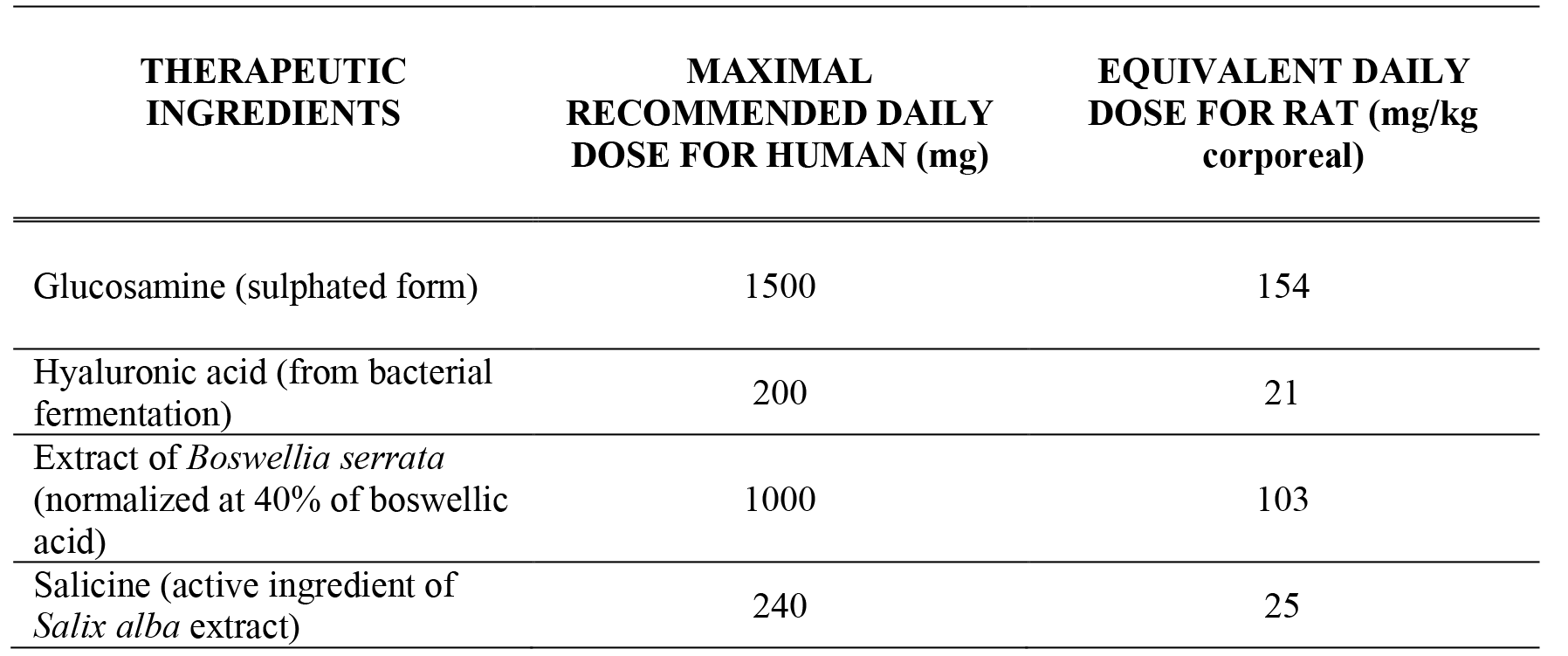
Daily doses of therapeutic ingredients recommended for human and equivalent doses for rat.

| THERAPEUTIC INGREDIENTS | MAXIMAL RECOMMENDED DAILY DOSE FOR HUMAN (mg) | EQUIVALENT DAILY DOSE FOR RAT (mg/kg corporeal) |
| --- | --- | --- |
| Glucosamine (sulphated form) | 1500 | 154 |
| Hyaluronic acid (from bacterial fermentation) | 200 | 21 |
| Extract of *Boswellia serrata* (normalized at 40% of boswellic acid) | 1000 | 103 |
| Salicine (active ingredient of *Salix alba* extract) | 240 | 25 |

Solutions of therapeutic ingredients were made daily as follows: GS (Novel Ingredients, NJ, USA), 200 mg/mL of water; HA from bacterial fermentation (A&A Pharmachem Inc, Ontario, Canada), 10 mg/mL of water; BS (40% of boswellic acid) (Dolcas Biotech, NJ, USA), 200 mg/mL of organic canola oil; SA (25% of salicin) (Novel Ingredients, NJ, USA), 50 mg/ml of water. Due to the specificity of BS’s solvent, all rats in groups not receiving treatment with BS received an equivalent daily volume of canola oil (100 μL). At the end of the experiment, all rats were euthanized by injection of a lethal dose of pentobarbital. A blood sample and knee cartilage of the two hind paws were collected. Plasma was extracted, samples were rapidly frozen by liquid nitrogen and preserved at −80 °C for future assays.

### Expression of Pro-Inflammatory Genes and Cartilage Degradation

Cartilage samples were reduced to powder with liquid nitrogen. RNA was extracted with *Pure LinkRNA Mini Kit* (Life Technologies, CA, USA; cat# MAN0000406, protocol with Trizol and DNase). Extraction purity was validated using spectrophotometry (absorbance ratio 260 / 280 nm). Inverse transcription was carried out on 400 ng of RNA for each extract according to *high capacity cDNA reverse transcription kit* method (Applied Biosystems, CA, USA; cat# 4368814). Obtained complementary DNA was used for real-time polymerase chain reaction essays (rt-PCR). Real-time PCR was performed with *SensiFAST SYBR No-ROX* kit from Bioline and with a LightCycler 480 from Roche (Mississauga, Canada). Three cytokines responsible for pro-inflammatory processes were targeted: interleukin-1 (IL-1β), interleukin-17 (IL-17A) and tumor necrosis factor (TNF-α). Primer sequences used for amplification are shown in Table 2. and were purchased from Sigma Aldrich (Oakville, Ontario, Canada). Gene expression was quantified by Cycle Treshold method (Ct). Amplification standard curve of each gene was performed and the specific amplification efficiency was verified with a minimal threshold of 1,8 (maximum 2). In order to standardize and compare the different assays, a pool of cDNAs of all groups was used as an internal calibrator. Gene quantification values were expressed relative to the gene quantification of two endogenous references, β-actin and glyceraldehyde 3-phosphate dehydrogenase (GAPDH). Ratio expressions for both references were similar; only results with the expression of β-actin are shown in this study. The specificity of PCR products was confirmed by migration on electrophoresis gel and melting curve analysis.

**Table 2.**
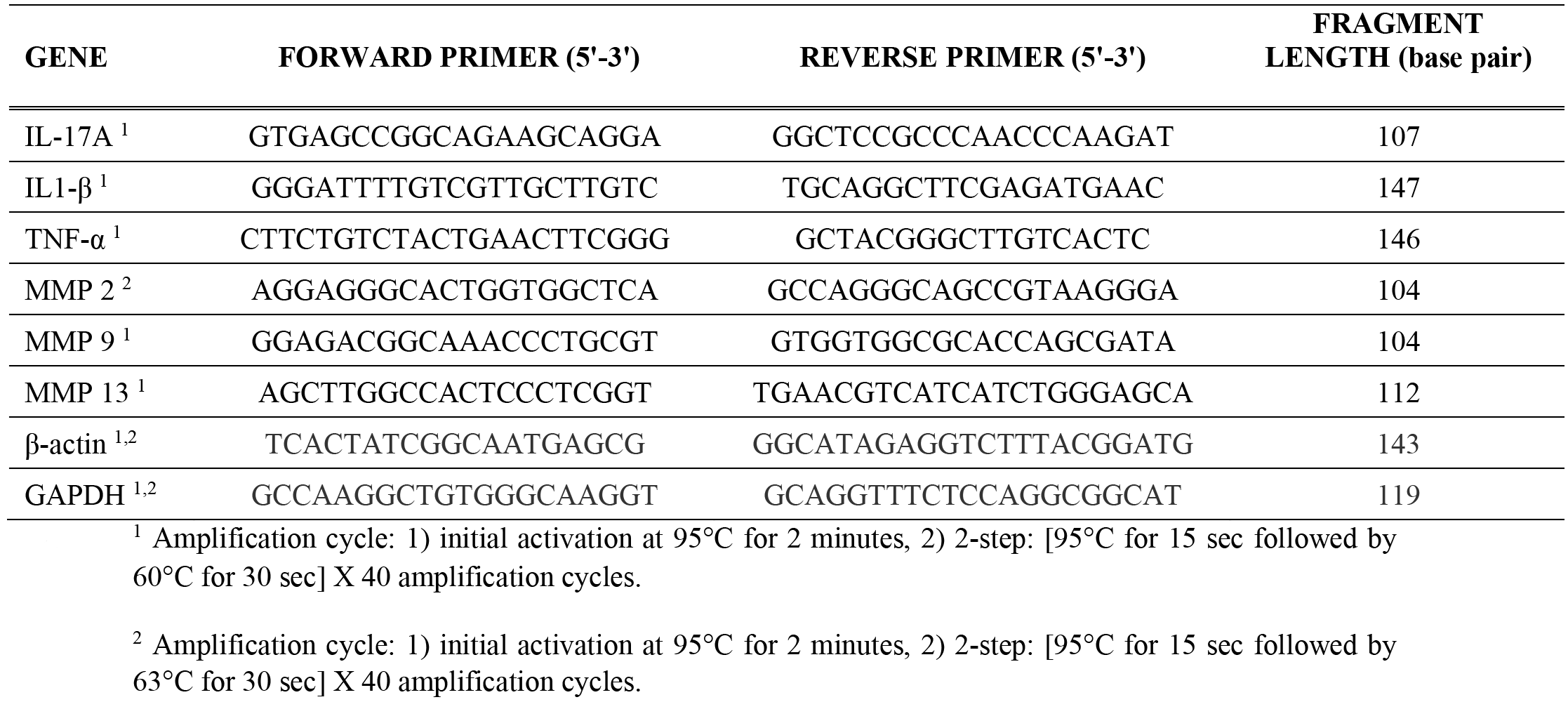
Primer sequences used for real-time PCR analysis.

### Total Antioxidant Activity

Total antioxidant activity of plasma was measured with the Antioxidant Assay Kit from Cayman Chemical (Ann Arbor, MI, USA; cat# 709001). Values were expressed in equivalent values of Trolox.

### Statistical Analysis

The results are shown as mean ± standard errors. They were analyzed using JMP Pro (SAS, Cary, NC, USA). One-way ANOVAs followed by a Tukey’s test were used to determine if there were differences between groups. The homogeneity of variance and the normality of data were tested using a Shapiro-Wilk and Bartlett’s test respectively. Two groups were statistically different if the p-value is lower than 0,05

## Results

### Effect on Clinical Signs of Arthritis

Measurement of hind-paw thickness and evaluation of the arthritis scores quantified the development of clinical symptoms in rats during the experiment. First signs of arthritis appeared on day 12 after the injection of Freund’s adjuvant (Fig. 2). On day 19, hind-paw thickness of the CFA group increased significantly (6.19 ± 0.90 mm) compared to the control group (3.59 ± 0.17 mm). Treatments with combinations of three ingredients (*Glu+Hyal A+Bosw*; *Glu+Hyal A+Sal*; *Glu+Bosw+Sal*) and *Glu+Bosw* limited articular swelling and significantly reduced hind-paw thickness during the treatment. At the end of the experiment (day 29), hind-paw thickness of these groups was significantly inferior to the CFA group (Fig. 2A) (*Glu+Hyal A+Sal*: 4.50 ± 0.78 mm; *Glu+Hyal A+Bosw*: 4.35 ± 0.65 mm; *Glu+Bosw*: 3.85 ± 0.63 mm; *Glu+Bosw+Sal*: 3.84 ± 0.48 mm; CFA: 5.84 ± 0.97 mm, p < 0.05). Severity of arthritis was evaluated by visual inspection through a score system which reflects the number of affected joints and the swelling intensity in digits and wrists/ankles. Arthritis scores increased from days 12 to 18 post-induction. At day 18, the maximal score was reached for the CFA group (Fig. 2B). A significant decrease of arthritis scores was observed for *Glu+Hyal A+Bosw*, *Glu+Bosw+Sal*, *Glu+Bosw* and *Glu+Hyal A+Sal* treatments in comparison to the CFA group at days 23, 25, 26 and 27 respectively. At day 29, arthritis scores of these groups were significantly inferior to the CFA group (*Glu+Hyal A+Bosw*: 5.33 ± 3.61; *Glu+Bosw+Sal*: 5.17 ± 4.07; *Glu+Bosw*: 5.17 ± 3.06; *Glu+Hyal A+Sal*: 5.83 ± 2.14; CFA: 11.42 ± 2.31, p < 0.01). Treatments with only glucosamine (*Glu*) and both glucosamine and hyaluronic acid (*Glu+Hyal A*) yielded no significant improvement of the clinical symptoms. The hind-paw conditions of each group at the end of the experiment are shown in Fig. 3..

**Fig 2.**
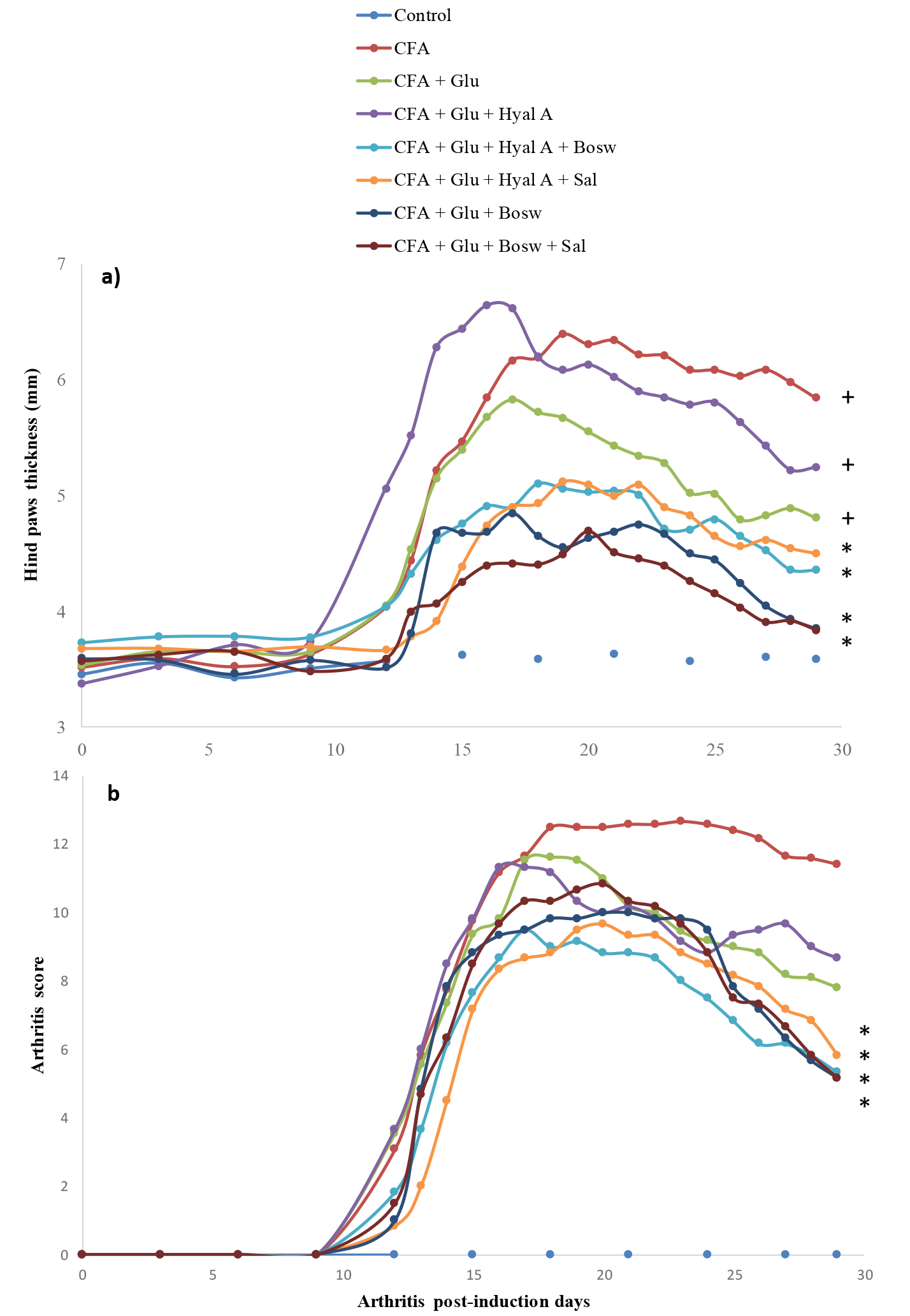
Effect of treatments on severity of arthritis. **a)** Hind paws thickness (mm) and **b)** arthritis score in time (number of arthritis post-induction days). Treatments have been daily administered starting at day 14 to day 29. Each circle is a mean ± SEM (control and CFA groups: n = 12; *Glu*: n = 11; *Glu+Hyal A*, *Glu+Hyal A+Bosw*, *Glu+Hyal A+Sal*, *Glu+Bosw* and *Glu+Bosw+Sal*: n = 6). Significant differences with CFA group (*) and control group (+) at day 29 are shown in **a)** p < 0.05 and in **b)** p < 0.01.

**Fig 3.**
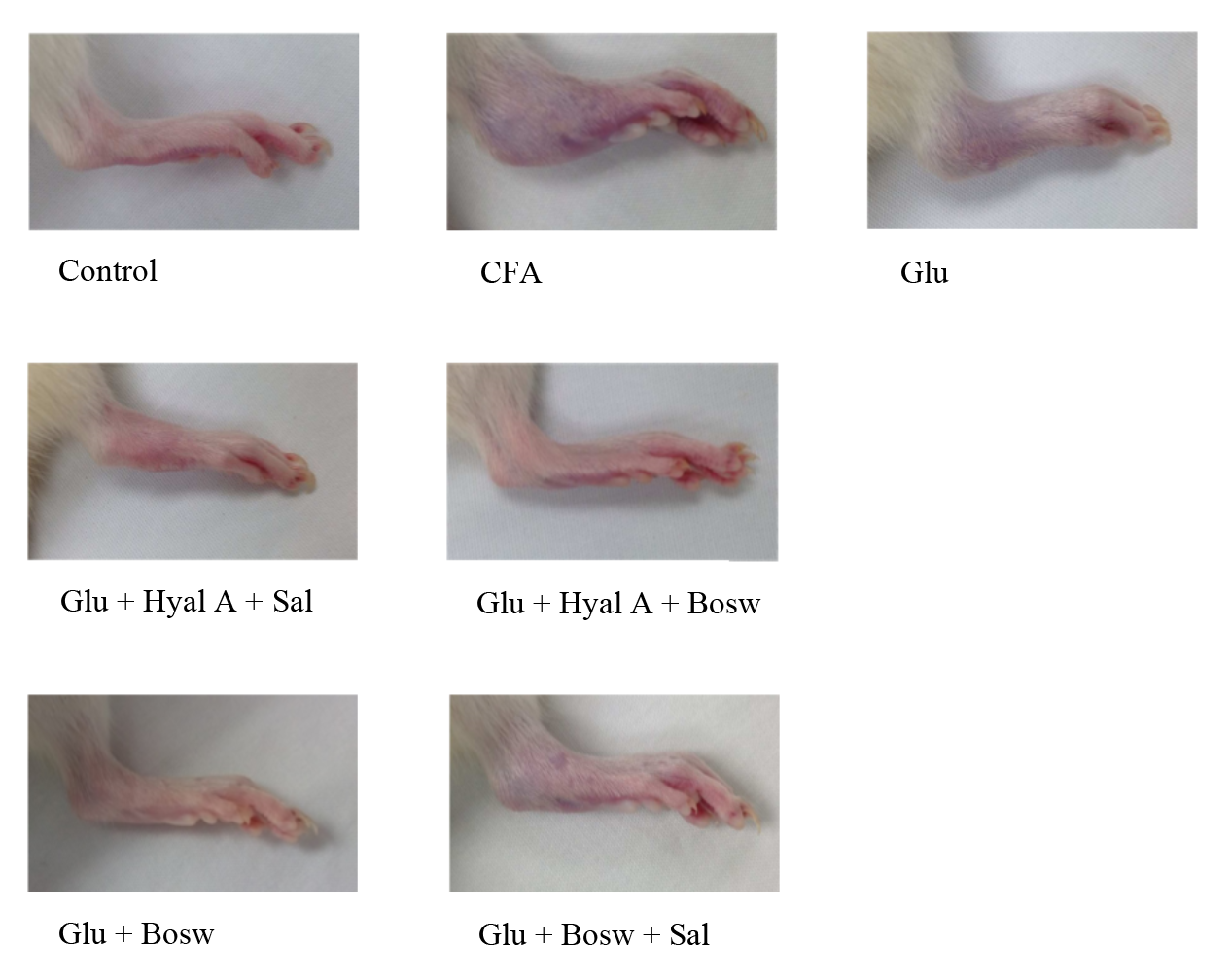
Clinical condition of hind paws at the end of the experiment (after treatment of 15 days).

### Effect on Expression of Pro-inflammatory and Cartilage Degradation Genes

At day 29, expression of cytokines *IL-17A* and *IL-1β* of CFA group was significantly greater than the control group (Fig. 4A-B). *Glu+Hyal A+Sal* treatment significantly reduced the expression of *IL-17A* and *IL-1β* compared to the CFA group. A decrease in the expression of *IL-17A* by *Glu+Bosw* and *Glu+Bosw+Sal* was also noticed. *Glu+Hyal A+Bosw* tended to limit *IL-17A* and *IL-1β* expressions, but the difference with the CFA group wasn’t significant. *TNF*-α expression of CFA group was not superior to the control group (Fig 4C). *Glu+Hyal A+Sal* treatment inhibited *TNF*-α compared to both the control and the CFA group.

**Fig 4.**
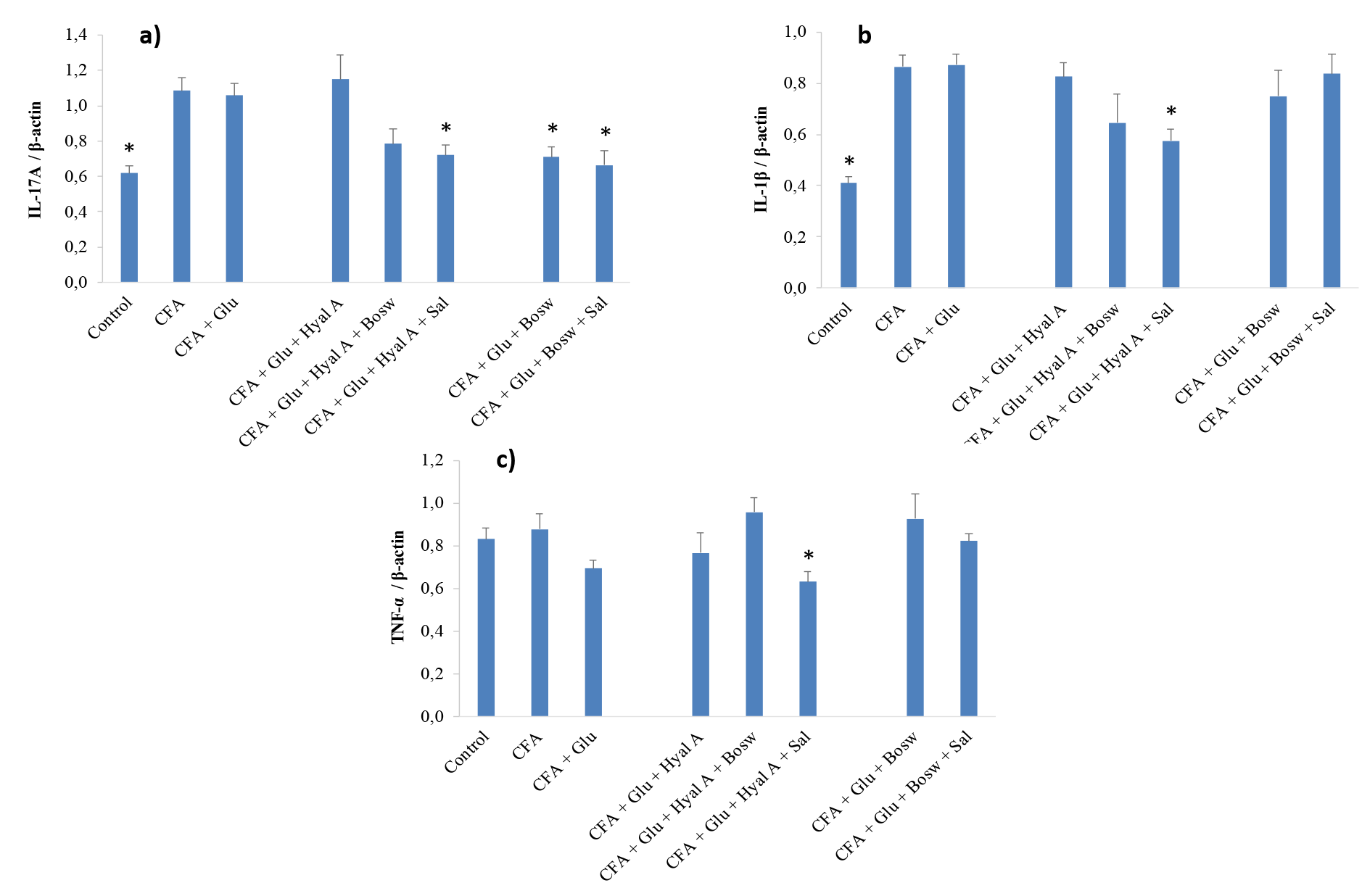
Effect of treatments on genes expression of pro-inflammatory cytokines. Genes expression of **a)** IL-17A, **b)** IL-1β and **c)** *TNF*-α were measured in articular cartilage of hind paws at day 29 (15e day of treatment). Each circle is a mean ± SEM (control and CFA groups: n = 10–12; *Glu*: n = 9–11; *Glu+Hyal A*, *Glu+Hyal A+Bosw*, *Glu+Hyal A+Sal*, *Glu+Bosw* and *Glu+Bosw+Sal*: n = 5–6). Significant differences with CFA group (*) at day 29 are shown (p < 0.05).

### Effect on Total Antioxidant Activity of Plasma

Fig. 5. shows that all treatments with combinations of two or three ingredients significantly increased the antioxidant capacity of blood, compared to the CFA group and control group (*Glu+Hyal A*: 4,16 ± 0,14; *Glu+Hyal A+Bosw*: 4,51 ± 0,07; *Glu+Hyal A+Sal*: 3,81 ± 0,18; *Glu+Bosw*: 4,71 ± 0,27; *Glu+Bosw+Sal*: 3,80 ± 0,06; CFA: 2,87 ± 0,16; Heal: 3,24 ± 0,13, p < 0,05).

**Fig 5.**
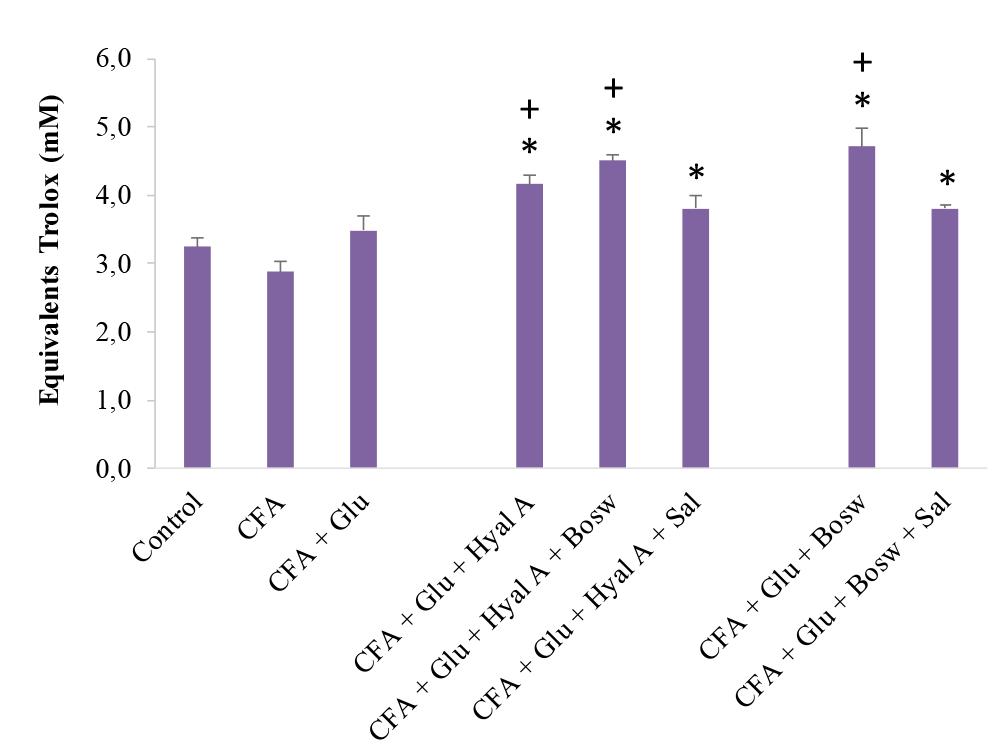
Effect of treatments on plasma content as total antioxidants. The content was measured in blood plasma at day 29 (15e day of treatment). Each circle is a mean ± SEM (control group: n = 8; CFA groups: n = 10–11; *Glu*: n = 9–10; *Glu+Hyal A*, *Glu+Hyal A+Bosw*, *Glu+Hyal A+Sal*, *Glu+Bosw* and *Glu+Bosw+Sal*: n = 5-6). Significant differences with CFA group (*) and control group (+) at day 29 are shown(p < 0.05).

## Discussion

The aim of this study was to evaluate if glucosamine sulfates therapeutic effects against rheumatoid arthritis could be enhanced through a combination with hyaluronic acid, *Boswellia serrata* extract or *Salix alba* extract. First symptoms of arthritis were evaluated by measuring the thickness of hind paws and by a visual examination (arthritis score). The combinations with BS and/or SA significantly reduce hind-paw thickness and arthritis scores during the treatment. *Salix nigra* bark methanol extract (100 mg/kg/day) has inhibited the progression of collagen-induced arthritis in rats at the end of the experiment by leaving arthritis scores and paw swelling close to healthy control (Sharma et al., 2011). In other studies, boswellic acid extract (total acid content: 93 ± 3%) from BS (250 mg/kg) was more efficient than glucosamine (250 mg/kg) to reduce inflammation in Mycobacterium-induced arthritis in acute and chronic model of inflammation in rats (Singh et al., 2007). Furthermore, in the same study, the combination of these two ingredients have shown a significant synergistic effect on chronic inflammation with a dose of 125 mg/kg for boswellic acid and 125 mg/kg for glucosamine. We suspect that the synergistic effect result from the combination of the different metabolic targets by which the bioactive molecules reduce inflammation. Anti-arthritic proprieties of each ingredient might have amplified the therapeutic effect of treatments. Glucosamine and hyaluronic acid have major structural roles in articular cartilage. Glucosamine stimulates the production of glycosaminoglycans that provide strength and elasticity to cartilage and connective tissues by holding joint tissue together and giving shock-absorbing properties (Singh et al., 2007). HA is a component of the synovial fluid and confers viscosity, as well as shock-absorbing and lubricating abilities (Moreland, 2003)., *Boswellia spp*. and *Salix spp*. do indeed produce active compounds like boswellic acid and salicin that show anti-inflammatory activities (Kimmatkar et al., 2003; Shara and Stohs, 2015). They directly target the inflammatory mediators such as interleukins and metalloproteinases (Umar et al., 2014; Sharma et al., 2011). BS extracts inhibit the 5-lipoxygenase which contributes to the progression of chronic inflammation through greater recruitment of white blood cells at inflammatory sites (Kimmatkar et al., 2003). BS and SA also have ROS-scavenging properties and can have a certain control on antioxidant enzymes (Sharma et al., 2011; Umar et al., 2014). Many studies, including ours, have concluded that combinations of glucosamine with either BS and/or SA are a promising strategy for limiting clinical signs of arthritis (Umar et al., 2014; Sharma et al., 2011; Kimmatkar et al., 2003). In our study, GS alone or in combination with HA did not result in any improvement of the arthritis symptoms. A previous study demonstrated that glucosamine can inhibit swelling in joints and reduce arthritic scores in rat adjuvant arthritis (Hua et al., 2005). However, the dose that they used was much higher than the equivalent dose usually administered to humans. It happens frequently in animal studies that the administered dose is higher than the recommended equivalent (when corrected for allometry) for humans. These doses could not be applied to humans according to the severe legislation in different countries. Maximal dosages of natural products are highly regulated to avoid adverse effects. We can therefore suspect that maximal recommended dosages for humans may not result in significant inhibition of the clinical signs of arthritis in both rats and humans. For example, a meta-analysis including 10 trials with an average of at least 100 patients concluded that glucosamine (1500 mg/kg/day: recommended dosage) was not effective against osteoarthritis, having no relevant clinical effect on pain or structure of affected joints (Wandel et al., 2010). Then, it becomes clearly advantageous to use combinations of ingredients to compensate for the small effect of a single ingredient (at recommended doses), and to rely on the synergetic effect of combinations without exceeding the recommended dosages of individual biomolecules.

Inhibition of pro-inflammatory cytokines can reduce clinical signs of arthritis. We therefore analyzed the mRNA content of three pro-inflammatory cytokines. IL-1β and TNF-α, two major pro-inflammatory cytokines, are both known to be present at high concentrations in serum and synovial fluid in patients with RA (Umar et al., 2014). IL-1β and TNF-α stimulate their own production and the production of other cytokines, amplifying the inflammation process (Moreland, 2003) and contribute synergistically to produce the inflammasome (Gaffen, 2009).

In our study, *TNF*-α and *IL-1β* expression was significantly reduced by only *Glu+Hyal A+Sal*. A previous study, on synovial-cell cultures from patients with RA, demonstrated that blocking the activity of TNF-α significantly reduced the production of IL-1, IL-6 and IL-8 (Butler et al., 1995). Thus, blocking TNF-α may have a greater overall effect on inflammation than only blocking IL-1. Moreover, we suspect that this reduction of *TNF*-α partly results from SA activity. In a recent review, Shara and Stohs, (2015) came to the conclusion that the anti-inflammatory activity of SA is associated with down regulation of the pro-inflammatory effect of TNF-α. In addition to salicin, SA has other active compounds like polyphenols and flavonoids which may also play a role in the therapeutic action of SA (Dragos et al., 2017). On the other hand, HA didn’t seem to have any impact on the level of *TNF*-α in RA rats. Injection of intra-articular HA in rat antigen-induced arthritis did show no significant changes in the level of TNF-α in short-term and in long-term experiments (Roth et al., 2005). Thus, we suspect that SA has an important role in the anti-inflammatory properties of a *Glu+Hyal A+Sal* treatment.

At the end of the experiment, three treatments containing BS and/or SA were able to inhibit *IL-17A* expression compared to *TNF-α* that was only inhibited by *Glu+Hyal A+Sal*. Furthermore, we did not observe a difference in the level of *TNF*-α expression between the control and CFA group. These results suggest that TNF-α had a weaker role in chronic inflammation than IL-17A, at least at the sampling periods of our experimentation. *IL-17A* is also over-expressed in RA (Dudler et al., 2000). It has been shown that TNF-α may be important in the onset of the arthritis induction, but it gradually loses its dominance with the progression of the inflammation (Joosten et al., 1996). They showed that anti-TNFα treatment was efficient shortly after the collagen-induced arthritis (CIA) in DBA/1 mice, reducing cartilage destruction, but that it had little effect when CIA is fully established (Joosten et al., 1996). To notice a difference in *TNF*-α levels, measurements should have then been performed at the beginning of the inflammatory phase. We suggest that TNF-α played a lesser role in the late phase of our experiment and had lower involvement in inflammatory modulation than IL-1β and IL-17A.

As mentioned previously, our treatments had a stronger impact on *IL-17A* expression than in the expression of the two other cytokines. IL-17 is involved in inflammation by stimulating other pro-inflammatory cytokines and metalloproteinases in synoviocytes and chondrocytes (Dudler et al., 2000). For example, it stimulates secretion of IL-1β and TNF-α by macrophages (Jovanovic et al., 1998). Two studies came to the conclusion that arthritis treatments involving inhibition of IL-17 could be as efficient as blocking IL-1 and TNF-α. They showed that IL-17 expression and activity is partly independent of these two cytokines under arthritis conditions (Koenders et al., 2005; Koenders et al., 2006). It can, for example, aggravate joint inflammation and cartilage destruction on its own without the increase of IL-1 or TNF-α. Also, blocking IL-1 in TNF-deficient mice was not sufficient to reduce IL-17 effects in streptococcal cell wall-induced (SCW) arthritis model (Koenders et al., 2006). IL-17 has the capacity to partly supplant the functions of IL-1 since these two have many overlapping responses and functions even if they are not from the same cytokine family, as shown in SCW-induced arthritis and IL-1-deficient mice (Koenders et al., 2005). We conclude that IL-17A might partly lead the inflammatory process at the end of our experimental arthritis and that treatments with plants were effective to decrease the expression of this cytokine. Anti-inflammatory properties of BS and SA successfully reduced the IL-17A effect according to our results.

Reactive oxygen species (ROS) can also contribute to matrix component degradation (Campo et al., 2008). ROS are generated at high rates in synovial neutrophils from RA patients (Sato et al., 1988). Synovial fluid and HA are respectively susceptible to degradation and depolymerization by a high level of ROS (Sato et al., 1988). These processes promote loss of viscosity in the joint as well as osteoclast activation (Filippin et al., 2008). Different antioxidants (polyphenols, tannins, etc.) are found in natural ingredients (Sato et al., 1988) which have been reported to partly protect and limit damage to cartilage (Venkatesha et al., 2011). All combinations with HA, BS and/or SA have significantly increased the total antioxidant level in the plasma of our experimental rats. Combinations without SA appear to have more impact on antioxidant levels than those with SA. In a previous study, it has been demonstrated that CIA in rats increases the activity of three important enzymes involved in oxidative stress management in plasma: superoxide dismutase, glutathione peroxidase and catalase (Sharma et al., 2011). They showed that these enzymes respond naturally to a higher concentration of ROS after arthritis induction, increasing their activities to protect tissues in the joint. The antioxidant properties of biomolecules in the present study may have helped to attenuate oxidative stress. Another study demonstrated that GS reduces superoxide radicals in a dose-concentration manner which partly explains its antioxidant activity (Xing et al., 2009). Synovial fluid and endogenous HA usually protect the articular tissues from oxidative damage. Excessive ROS decrease the HA content of articulation while addition of exogenous HA can decrease ROS levels in synovial cells of RA and buffer the impact on HA oxidation and decline (Sato et al., 1988). It has also been observed that extract of BS has improved the antioxidant level in CIA rat models, significantly decreasing ROS (Umar et al., 2014). *Salix nigra* bark methanol extract has contributed to attenuate oxidative stress in CIA rats (Sharma et al., 2011). BS and SA also has a phenolic compound showing antioxidant activities (Kokkiripati et al., 2011; Dragos et al., 2017). We suggest that a combination of antioxidant properties of these natural ingredients increases the antioxidant capacity in plasma of our experimental rats. We also suggest that this rise in antioxidant capacity can partly be responsible for the reduction of the clinical signs of arthritis.

## Conclusion

We conclude that the addition of *Boswellia serrata* and/or *Salix alba* attenuates clinical signs of rheumatoid arthritis in Freund’s complete adjuvant-induced arthritis in rats likely due to both their anti-inflammatory and antioxidant properties. Combinations with these plants have decreased hind-paw swelling and improved arthritis scores. Our results on clinical symptoms have been confirmed by some of our molecular markers. Treatments with BS and/or SA help to reduce inflammation and cartilage degradation by reducing effectively *IL-17A* expression and to a lesser extent, the expression of *IL-1β*. BS and SA have helped to create a redox status that might buffer oxidative stress through higher antioxidant capacity in plasma. This capacity may be partly responsible for the amelioration of clinical symptoms of arthritis.

## Acknowledgments

We are thankful to Samuel Fortin (Ph.D.) and Caroline Morin (Ph.D.) to their help for the familiarization with the arthritic model of Freund’s complete adjuvant.

## Funding

This work was supported by the Industrial Research Assistance Program (IRAP) of National Research Council Canada (NRCC).

